## Supplemental Table 1 for "Genetic variation in promoter region of the bovine LAP3 gene associated with estimated breeding values of milk production traits and clinical mastitis in dairy cattle"

| **Haplotype** | **g.37140485 C>G** | **g.37140513 T>C** | **g.37140514 A>G** | **g.37140644 C>T** | **g.37140681 G>A** | **g.37140767 C>T** | **g.37140789 T>G** | **Frequency** |
|---|---|---|---|---|---|---|---|---|
| H1 | **C** | **T** | **A** | **C** | **G** | **C** | **T** | **0.4272** |
| H2 | **C** | **T** | **G** | **C** | **G** | **T** | **T** | **0.1956** |
| H3 | **G** | **C** | **G** | **T** | **A** | **C** | **G** | **0.1668** |
| H4 | **C** | **T** | **G** | **C** | **G** | **C** | **T** | **0.1484** |
| H5 | C | T | A | C | G | C | G | 0.0109 |
| H6 | G | T | G | C | G | T | T | 0.009 |
| H7 | C | T | G | T | G | T | T | 0.0071 |
| H8 | G | T | G | C | G | C | T | 0.0069 |
| H9 | G | C | G | T | A | C | T | 0.0048 |
| H10 | C | C | G | C | G | C | T | 0.0047 |
| H11 | G | T | A | C | G | C | T | 0.003 |
| H12 | G | C | G | T | A | T | G | 0.003 |
| H13 | C | T | G | T | G | C | T | 0.0024 |
| H14 | C | T | A | C | A | C | T | 0.0024 |
| H15 | G | T | G | T | G | C | T | 0.0024 |
| H16 | C | T | G | C | G | C | G | 0.001 |
