## Supplemental Table 2 for "Genetic variation in promoter region of the bovine LAP3 gene associated with estimated breeding values of milk production traits and clinical mastitis in dairy cattle"

| **Loci** | **Genotype** | **LMY (kg)** | **305dMY (kg)** |
|---|---|---|---|
| rs720373055:T>C | TT (136) | 1.26$^{b}$ | -141.62$^{b}$ |
| | TC (61) | -136.50$^{c}$ | -83.61$^{b}$ |
| | CC (15) | 633.08$^{a}$ | 627.35$^{a}$ |
| | **P** | 0.00614 | 0.0030 |
| | **α** | -100.07 | -225.065 |
| rs720349928:G>A | GG (140) | 401.20$^{a}$ | 260.66$^{a}$ |
| | GA (59) | 288.78$^{ab}$ | 305.14$^{a}$ |
| | AA (13) | -192.14$^{b}$ | -163.69$^{b}$ |
| | **P** | 0.0001 | <.0001 |
| | **α** | 272.75 | 299.45 |
