## Supplemental Table 3 for "Genetic variation in promoter region of the bovine LAP3 gene associated with estimated breeding values of milk production traits and clinical mastitis in dairy cattle"

| Effect | | Wald chi square | Odds ratio | 95% CI |
|---|---|---|---|---|
| Breed*** | Sahiwal | 13.802 | 6.85 | 2.48-18.89 |
| | Karan Fries | - | - | - |
| P/calving** | 5 | 0.74 | 4.41 | 0.856-22.68 |
| | 6 | 1.76 | 4.61 | 1.78-11.95 |
| | 7 | - | - | - |
| S/calving* | 1 | 0.03 | 2.02 | 0.525-7.79 |
| | 2 | 8.58 | 8.87 | 1.602-49.17 |
| | 3 | 1.79 | 1.17 | 0.234-5.867 |
| | 4 | - | - | - |
